# Isotope-Free Mapping of protein:RNA Interactions Using fCRAC and trxtools

**DOI:** 10.64898/2026.05.19.726220

**Authors:** Nic Robertson, Jan Mikolajczyk, Andrea C. Garcia-Sandoval, Aleksandra Helwak, Maxeen Major, Anouk Emadali, David Tollervey, Tomasz W. Turowski

## Abstract

Defining high-confidence RNA interaction sites for specific proteins is essential to understand RNA biology, but existing methods face trade-offs between specificity, sensitivity, and experimental accessibility. Here, we present fluorescent <u>Cr</u>oss-linking and <u>a</u>nalysis of <u>c</u>DNAs (fCRAC), a mammalian-cell optimized update to the CRAC protocol. In fCRAC, a fluorescent adaptor is used, in place of radiolabeling, to visualize RNA-protein complexes during gel-purification. fCRAC retains the tandem affinity purification and stringent, denaturing conditions of classical CRAC, enabling nucleotide-resolution mapping of protein:RNA interactions with high signal to noise ratio. We initially tested fCRAC using RPP25L, a component the RNase MRP and RNase P complexes. RPP25L almost exclusively bound to predicted, single sites in the RNA components (RMRP and RPPH1), showing excellent selectivity with nucleotide resolution. To support analysis of UV cross-linking data for more complex targets, we developed the trxtools package and example pipeline for standardized processing, quality control, and analysis of data from fCRAC and related methods. We include tailored strategies for repetitive RNA classes, such as tRNA and rRNA, which can be challenging to analyze using other approaches. We applied fCRAC and trxtools to define the RNA interactome of human CYCLON/CCDC86, a nuclear protein previously implicated in oncogenesis. This revealed specific interactions with rRNA, tRNA and ncRNAs involved in pre-rRNA and pre-tRNA processing.

**Highlights:**

- Nucleotide-resolution definition of RNAs interacting with specific proteins, including rRNA and tRNA
- Stringent denaturing purifications and robust visualization steps, with no requirement for radioactive labelling
- Trxtools provides an integrated analysis pipeline with approaches for analyzing both single and multi-copy RNA species

## Introduction

RNA-binding proteins (RBPs) play critical roles in cellular biology through their associations with RNA^1^. Disruptions in these interactions are frequently implicated in human disease, and so defining the RNAs bound by a particular RBP is an important experimental goal^2^. However, the abundance of cellular RNAs can make uncovering biologically significant interaction targets challenging, as many proteins will non-specifically interact with abundant RNA species^3^.

This is particularly true when attempting to define interactions with ribosomal RNA (rRNA) and their precursors (pre-rRNA). The rRNAs account for 80% of total RNA, and non-specifically contaminate protein purifications unless stringent methods are used^4^. CRAC (<u>Cr</u>oss-linking and <u>a</u>nalysis of <u>c</u>DNA) was developed to meet this challenge, using tandem affinity purification including a denaturing, His-tag pulldown, to define interactions between RNA and specific proteins with nucleotide resolution^5,6^.

The original CRAC protocol used radiolabeling to visualize protein-RNA complexes purified from yeast cells. However, many labs are finding accessing radioactivity for biochemical experiments increasingly challenging, due to both cost and safety considerations. To address this, we developed an updated version of the CRAC protocol, fCRAC, that replaces radioactive signal from direct RNA phosphorylation with a fluorescent signal from the ligated 3’ linker. We also incorporate several optimizations to improve both yield and purity when performing fCRAC on mammalian cells, where starting material may be limited. Finally, we introduce a new analysis pipeline for CRAC data using the trxtools package, a set of bioinformatic tools for the analysis of transcriptional data. The example pipeline has been prepared as a GitHub repository using Snakemake^7^, to facilitate data sharing and reproducibility.

We initially tested fCRAC using RPP25L, a common component of RNase MRP and RNase P. Strikingly, we overwhelmingly recovered the expected interactions with the RNA components RMRP and RPPH1, with nucleotide precision. Then applied fCRAC and trxtools to map the complex RNA interactome of CYCLON. This nuclear protein is overexpressed in solid and haemotological malignancies, and was associated with disease progression and resistance to immunochemotherapy in Diffuse large B-Cell lymphoma (DLCBL)^8^. The fCRAC data identified substantial CYCLON binding to rRNA and the pre-rRNA processing factors SNORA73 (U17) and RMRP, as well as both tRNAs and RPPH1, which cleaves pre-tRNAs.

## Results

### fCRAC protocol overview

Recent progress in UV cross-linking methods revealed that commonly used CLIP-seq protocols can yield a substantial fraction of non-specific interactions due to the presence of other RNA-binding proteins during antibody-based immunoprecipitation (IP)^9^. The introduction of denaturing steps was a major advancement of the CRAC method, enabling the investigation of RNA-protein interactions involving highly abundant RNAs such as rRNA and tRNA. This strategy is maintained in the current fCRAC protocol.

To facilitate tandem affinity purification, fCRAC protocol utilizes a “bait” protein of interest, tagged with a tandem affinity purification tag **(Fig. 1A).** Tags can be attached to either C- or N-terminus of the protein. We recommend use of an N-terminal tag: FLAG-(DYKDDDDK)-Ala4-His8 (FH-N), or C-terminal tag: His8-Ala4-FLAG (C-HF)^10,11^. In each case the tag is only 20 aa in length, reducing the chances of adverse interactions with protein function. However, functional testing may still be advisable, if feasible. The tagged protein can be introduced by expression on a plasmid, as presented here using CYCLON as an example. Alternatively, cell lines can be CRISPR-edited so that the endogenous protein expresses the tag^12^. This option has the advantage that the experiment is then expected to report RNA interactions of the protein at its native expression level.

**Figure 1:**
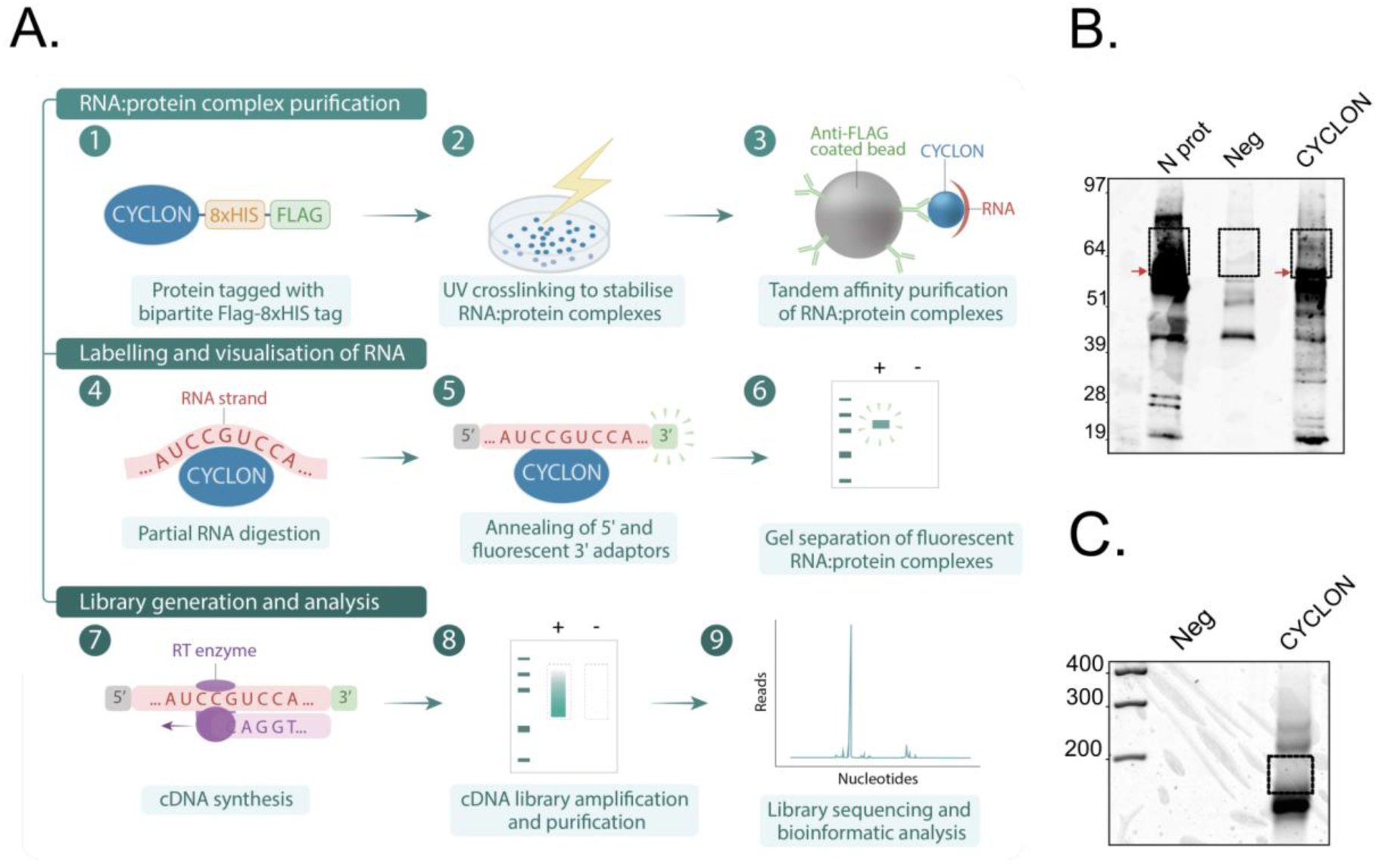
fCRAC and trxtools define the RNA interactors of bait proteins with high specificity, without the need for radioactive isotopes. **(A)** Schematic overview of the fCRAC method **(B)** Visualization of fluorescently labelled RNA co-purifying with FLAG-HIS tagged CYCLON protein. Protein:RNA complexes were resolved by SDS-PAGE, transferred into a nitrocellulose membrane and visualized via the fluorescent 3’ linker. Squares indicate the ideal target regions of the membrane to be cut and used for library generation. Ladder indicates molecular weight (kDa). “N prot”, nucleocapsid protein from SARS-CoV-2 was used as a positive control for cross-linking and protein:RNA complex purification. Red arrows indicated the expected position of uncrossed-linked protein. For CYLCON, the calculated molecular weight of FH tagged protein was ∼42 kDa, with uncrosslinked protein typically observed running at ∼60 kDa on test Western Blots. The N protein positive control is ∼48 kDa and observed to run at ∼55 kD. Two independent experiments were performed; the figure shows a representative result. **(C)** Visualization of cDNA libraries after PCR amplification. PCR products were resolved in a 3% MetaPhor Agarose gel. Squares indicate the regions that were excised and eluted for sequencing. Ladder indicates size of DNA fragments (base pairs). Two independent experiments were performed and a representative result shown.

In fCRAC, UVC (254 nm) cross-linking is used to stabilize RNA:protein interactions at sites of direct contact. Tandem affinity purification is then performed, first with an antibody against the FLAG-tag on the bait protein. An important step in the fCRAC protocol is protein footprinting, in which RNase is used to trim the RNA species bound to the purified protein. The aim is to have RNA lengths short enough to define closely interacting regions and be amenable to short-read sequencing, but long enough to allow efficient adapter ligation and confident read mapping. Different proteins protect bound RNAs to variable extents, so RNase treatment may need to be optimized for individual projects.

Following trimming, the RNases are robustly inactivated by addition of a strong protein denaturant, 6M guanidinium, and these conditions are maintained during initial binding and wash on the nickel column. While the RNA-protein complex remains bound to the nickel column, linkers (also termed adaptors) are added to both the 3’ and 5’ ends of the co-purified RNA. The 3’ linker is coupled with highly sensitive IRDye 800CW. This fluorescent moiety allows visualization of RNA after polyacrylamide gel electrophoresis, and optional RNA-protein transfer to membrane. The observed RNA-protein complexes are excised from the gel or membrane, and the protein components are digested with proteinase K. The RNA library is reverse transcribed, and PCR amplified ready for short-read sequencing.

The fCRAC protocol includes two points during which quality of the experiment can be visually assessed. First, following SDS-PAGE when specific signal from RNA bound by the bait protein can be compared to the negative control sample (**Fig. 1B**), in most cases a parental cell line which does not express tagged protein. A negative control should give a low level of background signal, whereas RNA co-purifying with bait protein should appear as a smear above the molecular weight of the non-crosslinked protein. As positive control, we used cross-linked cells expressing the N protein from SARS-CoV2, which avidly binds a diverse range of mRNA species^13^. We have released plasmids expressing codon-optimized versions of this protein (Addgene plasmid #157730)^10^.

Second, the final cDNA library can be visualized on an agarose gel **(Fig. 1C**). The library should appear as a smear above the adaptor-dimer band. We generally excise and purify the smear up to 200 bp, and a generate short-read library using 150 bp paired-end sequencing.

### Application of fCRAC to RPP25L

As an initial test of the specificity of fCRAC, we applied the method to determine the RPP25L interactome. Both RPP25L and its paralogue RPP25 are reported to be components of both the RNase MRP and RNase P complexes. However, published structures of both complexes have placed RPP25 but not RPP25L^13–16^. Both are RNP complexes, and the major binding partners of RPP25L were expected to be the RNA components (RMRP in RNase MRP and RPPH1 in RNase P; **Fig. 2A**).

**Figure 2:**
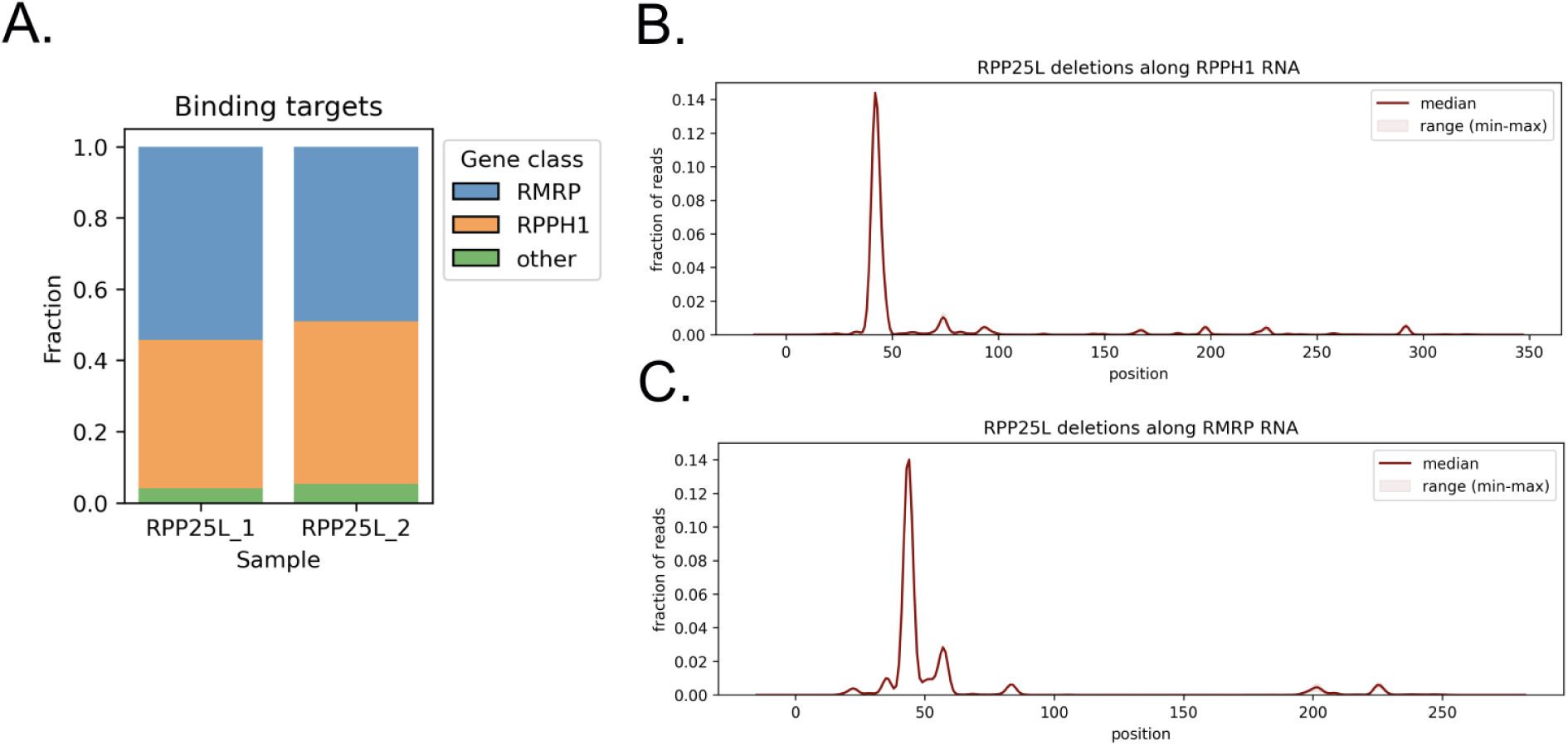
RPP25L almost exclusively interacts with the lncRNA components of the RNase MRP and RNase P complexes. **(A)** Binding targets of RPP25L in two biological replicates (individual CRISPR-tagged clones of K562 cells). **(B)** Deletions, indicating cross-linking sites, of the RNase P lncRNA RPPH1 with the RPP25L protein. **(C)** Deletions of RMRP (RNase MRP lncRNA) on RPP25L. Plots in B and C are representative of two experiments.

The genomic copies of RPP25L were tagged with a C-terminal HF-tag in K562, using a CRISPR-Cas9 approach^12^. fCRAC revealed that the vast majority of RNAs interacting with RPP25L were the known ncRNA components of the two complexes. RMRP averaged 48.8% of all mapped reads across two replicates, whereas RPPH1 averaged 40.8%. This indicates that fCRAC is highly specific with low levels of reads from contaminating non-specific RNAs including rRNA.

To further validate the method, we generated plots of single nucleotide deletion sites across RMRP and RPPH1 in the RPP25L cDNA sequence data. Reverse transcription, PCR amplification and Illumina sequencing can all introduce nucleotide substitutions, but nucleotide deletions are normally very rare. Deletions arise in fCRAC data due to reverse transcription errors at sites of direct crosslinking, where at least one amino acid remains associated with the RNA after protease digestion. This analysis demonstrated cross-linking sites between RPP25L and RMRP at positions +44, and with RPPH1 at positions +42 (**Fig. 2B and Fig. 2C).** These positions are different from the interaction sites for two other proteins in the RNase P and MRP complexes we previously mapped in RMRP and RPPH1 (POP1 and POP4; Robertson et al, 2022). We conclude that fCRAC can identify RNA:protein interactions at nucleotide resolution, and that RPP25L is an interactor of both RPPH1 and RMRP.

### Application of fCRAC and trxtools to CYCLON

Since RPP25L essentially bound only two sites in the entire transcriptome, little bioinformatic processing of the data is required. However, most RNA binding proteins do not show such simple interaction patterns. For these cases, we developed the trxtools package to accompany fCRAC data analysis. As a protein expected to show a more complex spectrum of interactions, which would require more sophisticated data analysis, we selected the human CYCLON protein. CYCLON is a nuclear protein localized to the nucleoplasm and nucleoli in DLBCL cell lines^18,19^ and in patient samples. The only characterized domain is shared with Cgr1 from yeast, which is implicated in nucleolar morphology and ribosome biogenesis^20^. Although CYCLON had been identified among RNA-associated proteins in high-throughput *in silico* and *in vitro* screens^21–25^, no information on RNA targets identity was available. To gain insight into its potential involvement in RNA metabolism or other nuclear processes, CYCLON-associated RNAs were identified using fCRAC and the resulting sequence data were analyzed with a new suite of tools, Trxtools.

### Data analysis with Trxtools

Analysis and interpretation of data from fCRAC and similar methods poses several bioinformatic challenges. Pipelines must be able to analyze canonical RNA classes that can be uniquely mapped (e.g. mRNAs and lncRNAs), as well as having dedicated strategies to handle highly repetitive loci such as rRNA and tRNA which are often excluded using other experimental approaches. Trxtools was developed to meet these requirements by providing a collection of functions and scripts for downstream analyses of RNA:protein interaction datasets. The framework offers multiple options for normalization, read aggregation, and metagene or window-based analyses, allowing flexible and in-depth exploration of diverse RNA-binding proteins. It also has built-in methods to generate visual outputs to assess the technical quality of experiments datasets and reproducibility between replicates.

To demonstrate this standardized and reproducible approach, we implemented an example end-to-end analysis pipeline based on trxtools using Snakemake and Jupyter notebooks, which ensures portability and scalability across datasets. Both trxtools and the accompanying example pipeline are distributed as a ready-to-use package via GitHub. This includes a standardized bioinformatics directory structure, and visualization notebooks. We also use a file-naming convention, removing the need for metafiles (for details, see **Supplemental File, sections 5 and 6**). While advanced programming skills are not required, effective use of the framework assumes a good understanding of bioinformatic concepts and basic computational skills.

### Trxtools utilizes Unique Molecular Identifiers

Our pipeline employs two approaches to data pre-processing to handle Unique Molecular Identifiers (UMIs, also called inline barcodes^26^) present in the 5′ linker, thereby improving data accuracy:

i. For standard applications, such as mRNAs and lincRNAs, UMIs are processed with dedicated bioinformatic tools (UMI-tools), to reduce PCR duplication artefacts. BigWig files that have undergone this deduplication step include “*umitools*” in their filenames and are located in a separate directory for easy identification.
ii. For repetitive and highly expressed gene loci, such as rRNA and tRNA, we use an alternative collapsing strategy that leverages UMIs but is less sensitive to sequencing errors. It involves collapsing raw reads followed by adapter removal. This approach is necessary because standard deduplication tools like UMI-tools tend to collapse reads with similar sequences, even when they are not identical, which can distort results in highly repetitive regions.

The pipeline generates BigWig tracks that can be visualized directly in the genome browsers, and gene expression count tables (featureCounts), that are useful for further downstream analysis. Our example repository contains notebooks designed for this purpose. We strongly recommend performing multi-step quality control of the data, as described below.

### Quality Control and Read Classes

<u>Notebook 0</u> (quality control and read classes) is employed to verify data integrity and reproducibility, as well as assist in troubleshooting challenging experiments. This analysis relies on gene expression count tables imported from featureCounts outputs^27^. We utilize a variety of quantification and normalization methods to calculate genome coverage metrics (**Fig. 3A-C).**

**Figure 3:**
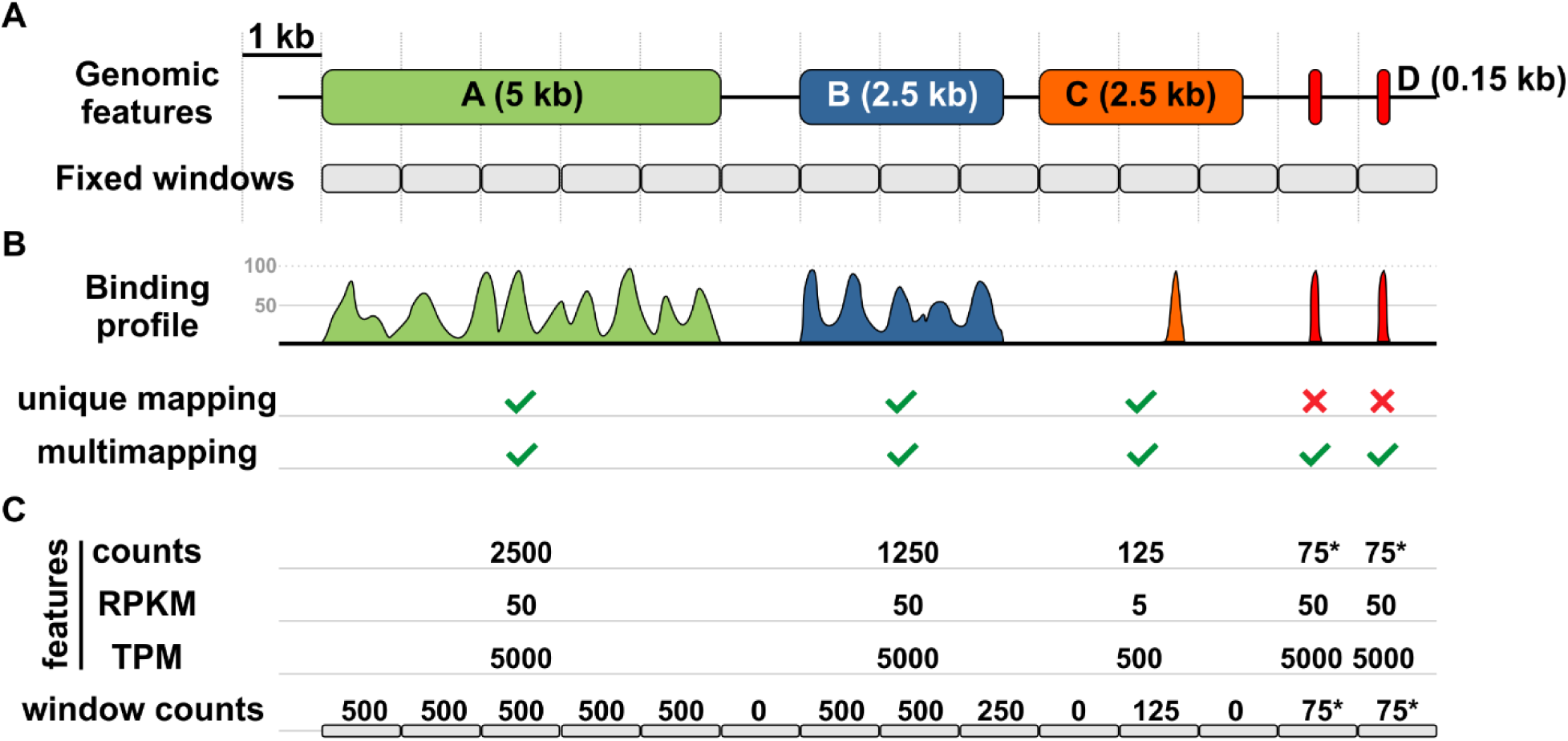
Analysis of data from fCRAC and related methods: quantification and normalization methods used by trxtools. **(A)** Genomic layout showing four genes of interest: A (5 kb, green), B (2.5 kb, blue), C (2.5 kb, orange), and D (0.15 kb, red). Region D contains a repeated element. Grey boxes represent annotation with fixed 1 kb windows. **(B)** Corresponding signal tracks (scaled 0–100) illustrating the distribution of signal intensity across each gene. Repetitive sequences are excluded from quantification based on uniquely mapped reads. **(C)** Summary showing differences in read counts obtained using commonly used normalization methods. Note that the lack of a normalization method leads to bias towards longer transcription units, whereas normalization strategies that take feature length into account (RPKM or TPM) bias towards shorter transcription units. Both strategies underestimate long transcription units with strong peaks (i.e. genomic feature C). The optimal strategy in such a case is to use fixed windows of a length closely related to the observed footprint size. Abbreviations: RPKM, reads per kilobase per million; TPM, transcripts per million.

The pipeline generates separate files for uniquely mapped reads and all reads allowing multimapping (unique + multi) **(Fig. 3B)**. In addition to raw feature counts, which represent the direct number of reads aligned to annotated genomic features, it also produces normalized counts in Transcripts Per Million mapped reads (TPM). This normalization method accounts for both gene length and sequencing depth (**Fig. 3C)**. RPKM is calculated by dividing read counts by gene length (kb) and by the total number of mapped reads (millions), thereby correcting for both transcript length and sequencing depth. TPM uses the same factors but normalizes for gene length first (reads per kilobase), followed by scaling across all genes so their values sum to one million, enabling direct comparison of transcript proportions between samples.

Despite the rapidly growing number of commonsense reasoning tools in artificial intelligence (AI) systems^28^, we still ultimately rely on human reasoning to assess the technical validity and biological significance of observed RNA:protein interactions. Two QC assessments are particularly important to guide this process. First, assessment of **read count (Fig. 4A)**. Trxtools generates summary tables and bar plots to display total read counts per library and mapped read counts per library. Negative controls should yield significantly fewer reads than experimental samples. High background or cross-contamination between samples should be suspected if this is not the case.

**Figure 4:**
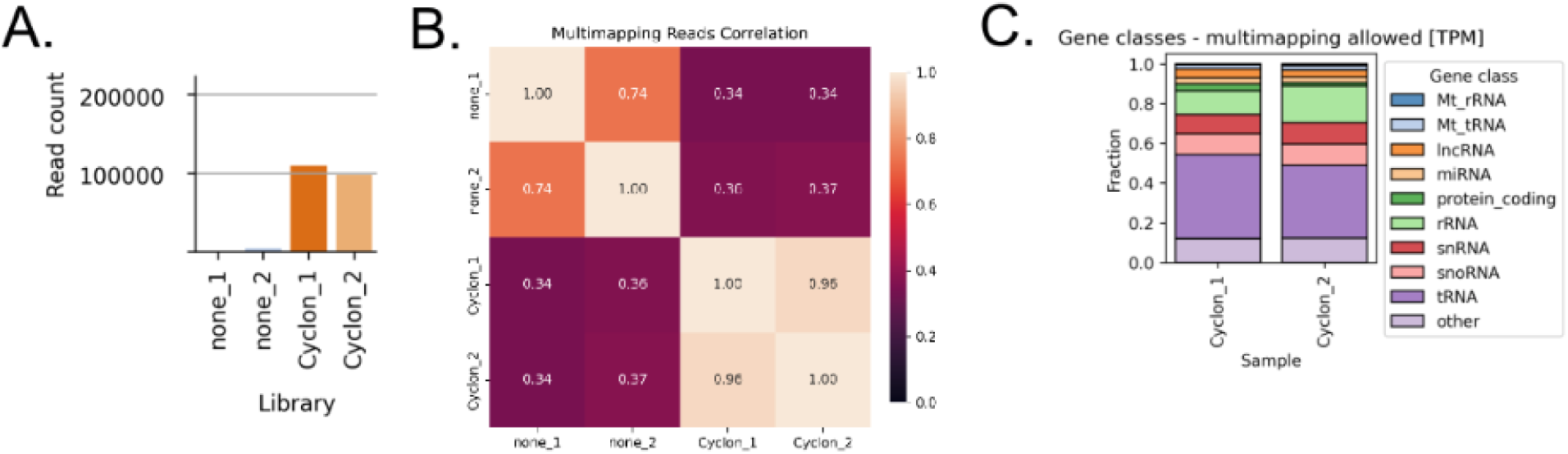
Trxtools provides an integrated analysis pipeline for fCRAC data including quality control assessments. **(A)** Mapped read counts (allowing multi-mappers) for example fCRAC data (from CYCLON and untagged negative control libraries). **(B)** Pearson correlations between CYCLON fCRAC TPM-normalized libraries based on multimapping reads table. **(C)** Gene classes of RNAs interacting with CYCLON (allowing multi-mappers), generated using Jupyter Notebook 0 (00_read_classes.ipynb). Abbreviations: Mt_rRNA: mitochondrial rRNA. Mt_tRNA: mitochondrial tRNA.

Second, **sample similarity and reproducibility** (**Fig. 4B).** Ideally, biological replicates should be more similar to each other than to other samples or negative controls. The example notebook provides visual measures to assess this, including principal component analysis (PCA) and correlation plots. These can be generated from three types of input data: raw feature counts for annotated genes, TPM-normalized data, and fixed-window tables using feature-agnostic, genome-wide annotations generated with a fixed window size (**Fig. 3)**. This last approach is more sensitive and less biased by expression levels and any background signal. It also offers better resolution, enabling more robust statistical analysis of binding sites and their reproducibility within features, which can be crucial if a feature, for example, has multiple binding sites. Fixed windows also facilitate comparisons within poorly annotated regions of the genome. The choice of the most suitable analysis depends on the specifics of the dataset and biological question.

Another analysis critical for both QC and further analyses is **gene class distribution**. It is represented using stacked bar plots that can be generated separately for uniquely mapped and multimapping reads and can use either raw counts or TPM-normalized values **(Fig. 4C).** For clarity, less abundant classes may be grouped into an “Other” category. Mapping to read classes enables more biology-driven QC. Similarity of class distribution between samples and the negative control is frequently observed in protein profiling data and is generally accepted, provided there are significantly more sample reads than in the negative control. This is because there will always be some degree of cross-contamination from samples processed in parallel and/or from barcode misreading.

Detailed analysis for different gene classes is outputted in separate notebooks. First, <u>Notebook 1</u> (01_ mRNA_lncRNA_snoRNA_snRNA.ipynb) contains templates for the standard subjects in the CLIP/CRAC analysis, namely mRNA, lncRNA, snoRNA, and snRNA. These are relatively easy to analyze, mapping uniquely to the genome and mostly at lower abundance with no expected saturation of UMIs. For these gene classes, trxtools uses the most stringent preprocessing criteria (UMItools and unique mapping), which efficiently eliminate PCR artefacts (visible as “towers” – groups of sequences with identical endpoints) that can otherwise bias results.

Second, <u>Notebooks 2</u> (02_human_rDNA.ipynb) and <u>Notebook 3</u> (03_human_tRNA.ipynb) contain results for rRNA and tRNA, respectively. For both classes, the pipeline uses a less stringent deduplication strategy and includes multimapping reads. Collapsing these reads with mRNA-dedicated tools introduces artefacts due to shared sequence fragments.

### Metagene binding profiles

Metagene binding profiles are a powerful way to visualize binding patterns and explore hypothesis about the biological function of particular RBP and trxtools provide a convenient way to create a variety of different profiles (**Fig. 5**). To generate metagene profiles, read coverages uploaded from BigWig files are aligned in a strand-aware manner relative to one of the predefined genomic features provided as BED-like annotation tables. This generates feature-level signal matrices (trxtools function: *getMultipleMatrices()*). Features could include: transcription start sites (TSS), annotated 5′ ends of ORFs, exon–intron junctions, 3′ ends of ORFs, and annotated 3′ ends of mRNAs. Read coverage is aggregated with respect to these feature coordinates producing average metagene profiles to assess binding enrichment patterns (**Fig. 5C, panels i and ii**). By default, aggregating features into a metaprofile uses averaging, but the *metaprofile()* function also allows using the median or the sum.

**Figure 5.**
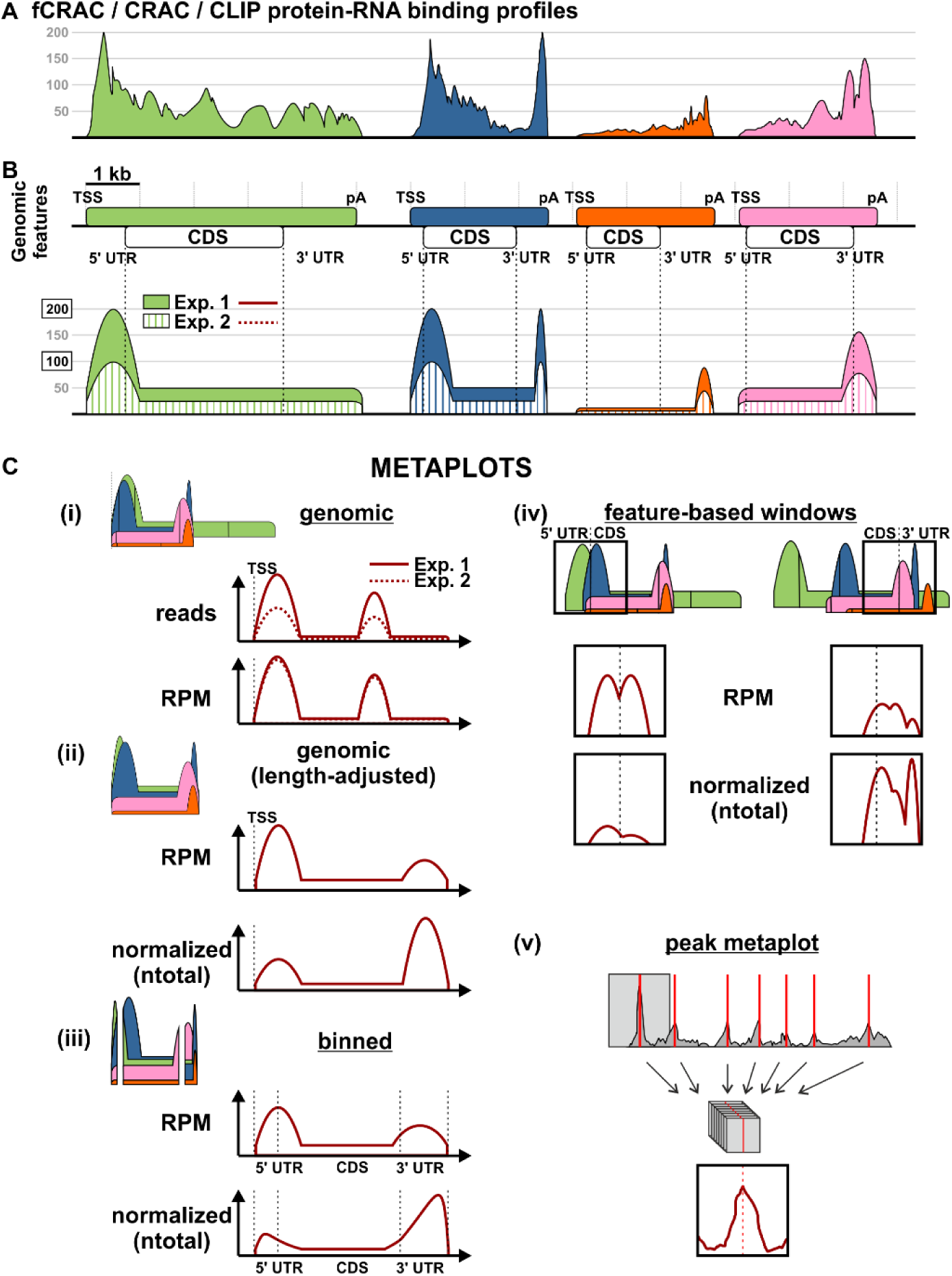
Illustration of normalization strategies and biases introduced by signal aggregation. **(A)** Representative signal profiles across four genes of different lengths and expression patterns (green, blue, orange, and pink). Profiles are characteristic of fCRAC signals generated by RNA-binding proteins (scaled 0–200). **(B)** Simplified coverage profiles illustrate read density across each gene. Two datasets with different sequencing depths are shown to demonstrate the significance of RPM normalization (see **C, panel i**). The plots show signal distribution, including enrichment at gene feature boundaries and local peaks. **(C)** Metaplots showing the consequences of different quantification and aggregation strategies. Aggregation can be performed using the mean, median, or sum. **(i) Genomic.** Superposition of signals from multiple genes aligned at the transcription start site, illustrating how genes of different lengths contribute unevenly to the combined profile. In contrast to raw read counts, RPM normalization highlights reproducibility of binding patterns. **(ii) Genomic, length-normalized.** Superposition of signals demonstrating bias toward regions with higher coverage or shorter genes. RPM normalization is contrasted with ntotal normalization, in which each gene or feature contributes equally to the metaplot. **(iii) Binned.** Each gene is divided into an equal number of bins. Multiple features can be incorporated into the binning strategy to quantify enrichment across defined regions. **(iv) Feature-based.** Windows centered on selected, pre-annotated features are aggregated to generate signal distribution patterns. Two examples are shown: start codon and stop codon. **(v) Peak metaplot.** Aggregation of signals centered on individual features (e.g. peaks or troughs) across gene or multiple genes. In *ntotal* normalization, each gene or feature contributes equally to the resulting metaplot. Abbreviations: TSS, transcription start site; pA, polyadenylation site; UTR, untranslated region; CDS, coding sequence; RPM, reads per million; TPM, transcripts per million.

Additionally, metagene binding profiles might be computed using a binning approach (trxtools function: *binMultipleMatrices()*, **Fig. 5C, panel iii**). Each gene is divided into an equal number of bins, and reads are aggregated within these bins. This method allows for direct comparison of binding profiles across genes of different lengths. A similar approach is applied to transcript elements such as untranslated regions (UTRs) and coding sequences (CDS).

While metaplots are a powerful tool, they should always be interpreted alongside other data to avoid overgeneralization or misinterpretation. Therefore, trxtools introduces a normalization feature (internal or so-called “ntotal” normalization). In this approach, each feature-level profile (e.g. each gene or each binding site) is independently scaled by its total signal, ensuring that all features contribute equally to the resulting metaprofile (**Fig. 5C, panel ii)**. This prevents the aggregate profile from being dominated by a small number of highly expressed genes and emphasizes relative positional enrichment patterns along aligned features rather than absolute read counts. Because this normalization relies on within-feature signal distribution, features with insufficient or sparse coverage should be excluded prior to aggregation to avoid introducing noise or artifacts.

The availability and accuracy of metagene alignments depend critically on the quality of the underlying genome annotation, which can vary substantially between species and annotation versions. In particular, incomplete or imprecise annotation of TSS location and UTRs may limit the interpretability of feature-centered binding profiles. Our notebooks also support data filtering, which can be used to focus on specific gene clusters provided by an external source, such as a selected GO term or genes falling into a selected quantile of expression

Feature-based metaplots typically use pre-annotated features such as the start codon in a CDS or the beginning of the 3’ UTR (**Fig. 5C, panel iv**). Importantly, these features can be expanded by using calculated features or additional datasets. Calculated features may represent DNA GC content or RNA:DNA hybrid energy, whereas additional datasets may provide such information as nucleosome positions or RNA m^6^A modifications. Moreover, one dataset may be used to produce features for the analysis of another dataset. A particular case of features is peaks or troughs detected in profiles (i.e. using calculated features or other datasets, **Fig. 5C, panel v**). This type of analysis provides data for comparing correlations between features and datasets, allowing testing of new biological hypotheses.

### Analysis of Data with Repeated Elements (rRNA and tRNA)

rRNA and tRNA are less typical targets of bioinformatic analyses due to the prevalence of non-specific signals in many datasets. However, the high signal-to-noise ratio achieved by fCRAC enables meaningful analysis of these species. Interpretation should still account for potential non-specific binding and the naturally high abundance of these transcripts in input samples.

rRNA genes have a complex structure, and accurate analysis can provide insights into protein-binding dynamics. The human rDNA transcription unit produces a long polycistronic transcript containing 18S, 5.8S, and 28S regions separated by external and internal transcribed spacers (ETS and ITS), which include processing sites^29^. Although multiple transcription units exist, genome sequence files contain only limited copies (within the human hg38 genome, 3 copies are included within the reference chromosomes, all located on chromosome 21), which simplifies analysis. Moreover, not all genome annotation files include all rDNA genes (most include only 5S rRNA). Therefore, we uploaded custom human rDNA annotations to the example repository: three transcription units from chromosome 21 - RNA45SN1, RNA45SN2, and RNA45SN3, differing in element lengths except for 18S and 5.8S.

Our <u>Notebook 2</u> includes tools to generate plots of all human rDNA sequences, marking the boundaries of individual elements and enabling comparisons across datasets (**Fig. 6A**). Additionally, 5S rRNA is encoded by multiple genes and pseudogenes distributed throughout the genome (e.g., RNA5S1, RNA5S2). For simplicity, we provide a dedicated metaplot analysis for 5S rDNA (**Fig. 6B**).

**Figure 6:**
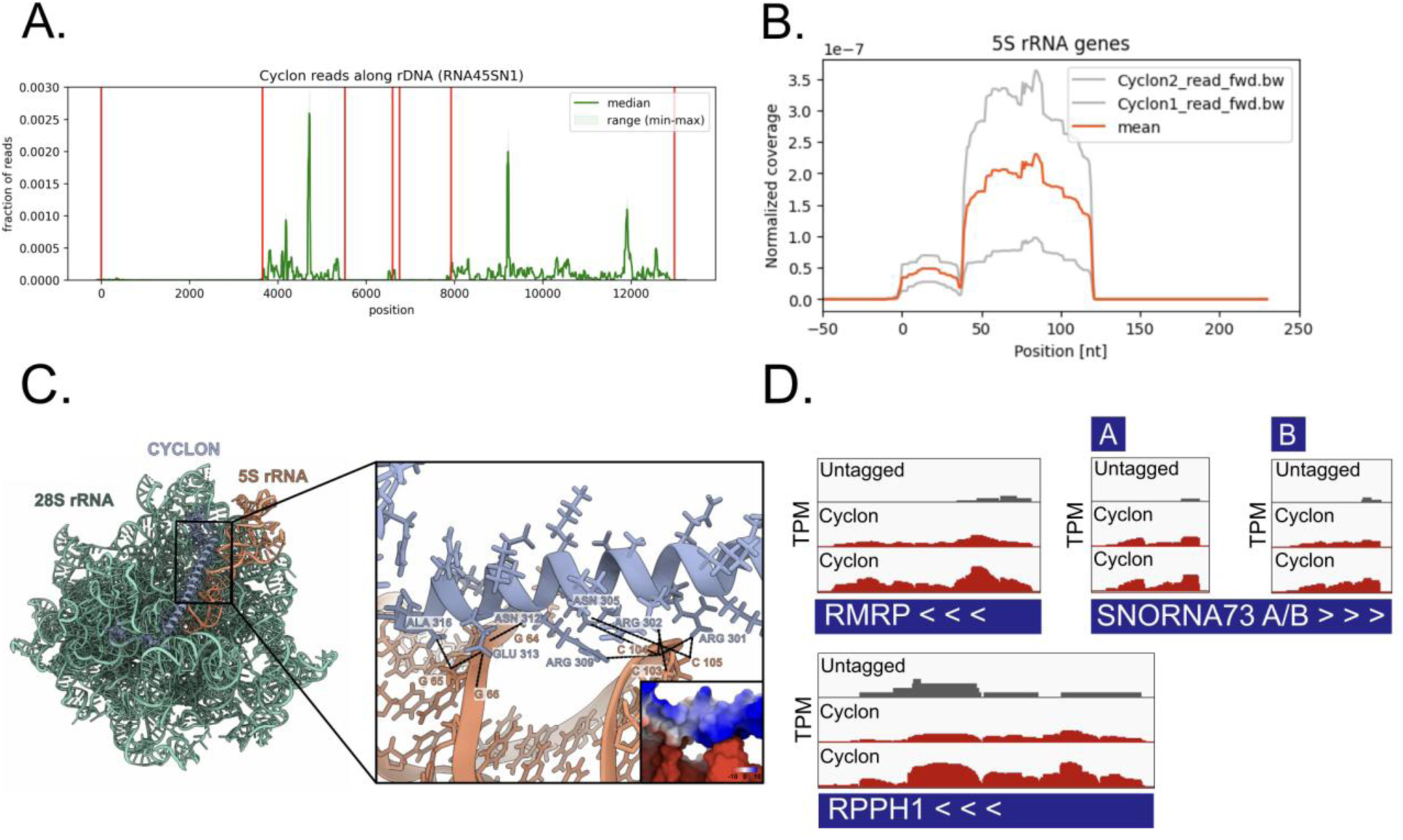
RNA interactors of CYCLON, as defined by fCRAC and trxtools. **(A)** rRNA and pre-rRNA reads from CYCLON fCRAC, plotted across RNA45S. Representative of results from two independent experiments (preprint doi: 10.64898/2026.05.13.724782) **(B)** 5S rRNA reads, averaged across different transcripts. Representative of result from two independent experiments. **(C)** Structure of CYCLON is pre-60S particles, showing interactions between CYCLON and the 5S rRNA. Structure PDB:8FL3^33^. **(D)** IGV plots of normalized read counts (transcripts per million, TPM) for selected RNA species highly enriched in CYCLON fCRAC. Plots show two independent experiments.

tRNA analysis, covered in <u>Notebook 3</u>, uses the same input files as the rRNA analysis (with less stringent collapsing), but it applies a similar methodology and visualization strategy to that for mRNA. A common challenge is the absence of tDNA annotations in standard annotation files. This can be resolved by downloading a gtf file with tDNA features from the gtRNAdb database^30^ (https://gtrnadb.org/). Binding patterns across tDNA are analyzed at the 5′ and 3′ ends, and additional features, such as terminator sequences, can be extracted from the sequence using custom Python functions and used as annotations.

### Characterization of RNA interactors of CYCLON

Plasmid-expressed CYCLON-His/FLAG (HF) was localized in the nucleoplasm and nucleoli of HeLa cells using immunofluorescence with anti-FLAG antibodies (**Fig. S1A).** Following anti-FLAG pulldown, the tagged protein was detectable by immunoblotting (**Fig. S1B**). Crosslinked CYCLON-HF was strongly enriched for copurified RNA, visualized by ligation of the fluorescent linker, relative to negative control samples (**Fig. 1B**). cDNA libraries were generated from CYCLON-bound RNAs by RT-PCR, and products in the expected range (100-200 bp) were visible following gel electrophoresis (**Fig. 1C**).

Sequencing of two fCRAC cDNA libraries yielded high-quality data as confirmed by the trxtools quality control metrics. The untagged negative control yielded more 20-fold fewer reads than CYCLON libraries (**Fig. 4A)**. There was good correlation between replicates (Pearson correlation *0.92*) (**Fig. 4B**), and negative controls and true samples clustered separately, indicating high reproducibility and low levels of cross-contamination.

Since other data indicated CYCLON’s role in ribosome biogenesis (preprint doi: 10.64898/2026.05.13.724782), our analysis aimed to include all RNA species, including rRNA. We initially selected multi-mapping as the most appropriate quantification method. Mapping of CYCLON reads initially revealed strong binding to regions annotated to protein-coding genes (for plots, please refer to Notebook 0). However, this high total coverage was a result of sparse reads identified by IGV inspection at many dispersed positions across the lengths of mRNA introns. To exclude this artefact, we performed analysis of RNA classes using TPM-normalized reads. This reduced the significance of very long features with sparse reads, which did not present confident protein-RNA interactions.

After this normalization, overall, the largest RNA class was tRNA (31.9%, averaged across two replicates), followed by rRNA (17.8%) and snoRNA (15.8%, **Fig. 4C)**. snRNA (10.1% of reads) and lncRNA (5.4%) were also highly represented. TPM-normalization is expected to boost the representation of tRNA, given their short size (typically 76-90 nucleotides^31^). However, without the normalization, tRNA still accounted for ∼10% of reads, supporting the conclusion of significant enrichment **(Fig. S2A**). tRNA are transcribed by RNA polymerase III (Pol III) and undergo multiple processing steps including cleavage of 5’ leader sequence by RNase P^32^. Notably, CYCLON also interacted with the catalytic RNA component of RNase P, RPPH1 (ranked 11 in uniquely mapping interactors, with 1.00% of reads), implying that CYCLON may have an under-explored role in tRNA metabolism. Supporting this, normalized tRNA metaprofiles aligned to both 5’ and 3’ end of the RNA showed that some of the recovered tRNA were pre-tRNA, e.g with the 3’ of 5’ leaders still intact **(Fig. S2B)**. We also note that mature tRNAs are normally strongly under-represented in RT-PCR based sequencing, due to their extensive modifications, supporting interactions with pre-tRNAs.

The observed rRNA interactions were further investigated by plotting reads across the polycistronic 45S transcript (**Fig. 6A).** Signals within the mature rRNA were visible as peaks in the 18S rRNA (around position 4900 in the 45S pre-rRNA) and in the 28S rRNA (around positions 9400 and 12,100 in the 45S pre-rRNA). Additionally, we prepared profiles for 5S rRNA, averaging across the multiple genomic copies of this rRNA. This revealed a major peak around 80–100 nt, accompanied by a corresponding peak in deletion frequency around nucleotide +81 **(Fig. 6B**).

To validate these results, we compared them with available structural data for CYCLON in the human nuclear pre-60S ribosomal particles, where CYCLON incorporation coincides with that of the 5S ribonucleoprotein (PDB: 8IPX-Y^25^ and PDB: 8FL2-4^26^). We selected structure 8FL3 (State I2) because it has the best resolution and harbors most of the interactions between CYCLON and rRNA. In this structure, CYCLON contacts the 5S at positions 103–105 bp, with an additional interaction at 65 bp, in good agreement with our findings (**Fig. 6C**). In contrast, the CYCLON fCRAC deletion peaks in 28S (peaks at positions 9210, 10483 and 11906 in 45S) are not clearly recapitulated by the reported CYCLON:28S interaction sites, both in terms of nucleotide number and spatial position in the structure. This may be partly due to the incomplete modeling of these rRNA regions, but could also suggest that the fCRAC deletion peaks may be coming from a different intermediate particle that has not yet been structurally captured.

Inspecting uniquely-mapped data showed that the overall most enriched individual transcript was RMRP, accounting for 4.3% of uniquely-mapped reads. RMRP is evolutionarily related to RPPH1, and is the lncRNA core of the RNase MRP complex that cleaves pre-rRNA in at least one location in humans (**Fig. 6D)**^12,34^.

In conjunction with the enrichment of rRNA and tRNA, these results suggest that CYCLON is part of an RNA processing hub. Other hits supporting this model include SNORA73 (U17), a box H/ACA snoRNA that interacts with the 18S and 28S region of the pre-RNA and is required for processing of the 5’ETS region^35^. Both paralogues (SNORNA73A and SNORNA73B) of this snoRNA were enriched: SNORNA73B was ranked 3rd in individual hits with 1.76% of reads, while SNORNA73B ranked 15th with 0.91% (**Fig. 6D)**.

In summary, these data first demonstrate that CYCLON specifically interacts mainly with tRNA and rRNA, and with the RNA components of the complexes that process these species.

## Discussion

Numerous methods have been reported, utilizing in vivo UV crosslinking combined with immunoprecipitation with specific antibodies. These are collectively known as cross-linking and immunoprecipitation (CLIP) and related terms, and they capture native proteins along with their interacting RNAs^36^. These techniques allow analysis of the native protein, without additional modification, but generally have significant background signals from abundant RNA species, since immunoprecipitation cannot be performed under strongly denaturing conditions. Tandem affinity purification (TAP) was initially used for protein complex isolation, and has long been recognized as giving much greater enrichment, and generally higher yield, than simple immunoprecipitation^37^. The original CRAC protocol took advantage of tandem purification but included a poly-His motif within the TAP tag (in place of the original calmodulin-binding peptide (CBP)) to allow a protocol that includes a highly denaturing step. In the fCRAC protocol, a 8xHis element is retained, but is linked to only a single FLAG peptide, generating tag of only 20 aa^11,38^.

Other protocols that allow high-stringency, denaturing purification have been developed; e.g. CLAP (Covalent Linkage & Affinity Purification)^9^. However, the CLAP protocol lacks a step to allow visualization, and so validation and gel enrichment, of the co-purified RNA before sequencing. In the original CRAC protocol, visualization was mediated by 5’-^[32]^P radiolabeling of the RNA molecules attached to the tagged protein. This has great sensitivity, but added a step to the protocol, while radiochemicals are expensive and their use is being increasingly restricted in many locations. We therefore included a fluorescent label in the 3’ linker, to follow the protein-bound RNA species. As the linker ligation is required in any event, this minimizes the additional work required.

Analysis of CRAC/CLIP data remains challenging due to the highly variable natures of these datasets. Standard tools developed for RNA-seq analysis or for detecting chromatin-binding features do not adequately address the specific requirements of these analyses. Feature-independent enrichment analysis has shown promise and has been applied previously^39^. However, this approach still lacks comprehensive statistical frameworks. As a result, many analytical decisions must be made by researchers, based on laborious and often subjective quality-control procedures.

Here, we demonstrate that fCRAC plus trxtools can provide highly specific information on the RNA interactors of mammalian proteins, without the need for radioactivity or specialized equipment. The analysis of RPP25L demonstrated the high signal to noise ratio of fCRAC: 90% of all mapped reads were recovered in the RNA components of RNase P and MRP with minimal recovery of the very abundant rRNAs and tRNAs. RPP25L is not visible in current structures of these complexes, which instead include its paralogue RPP25^14,15^. Future structural studies could aim to determine when and where RPP25L is incorporated.

On RNA-binding proteins with more complex patterns of interaction - which is most species - potential problems arise in analyzing fCRAC and related data, stemming from difficulties normalizing data from multi-mapping RNAs. To address this, trxtools provides an easy-to-use package to produce robust analysis and quality control metrics that help avoid common issues. The flexibility of trxtools particularly aids exploratory data analysis, to help draw meaningful conclusions from complex data. Applying this to CYCLON allowed the attribution of a large number of binding sites across a wide spectrum of RNA biotypes.

## Limitations

The fCRAC method does have some limitations. Most importantly, it works best when starting with a relatively large quantity of input material. We typically use 25-50 million cells per sample, and more might be required for proteins with very low expression levels. Future optimization will aim to reduce this requirement, to enable fCRAC to be used more widely in samples with limited availability.

Both trxtools and the example pipeline require familiarity with command-line syntax and a basic understanding of Python. Moreover, performing the analysis requires a solid grasp of core bioinformatics concepts. This may involve a learning curve, particularly for early-career researchers. Nevertheless, we have attempted to provide comprehensive considerations and guidance for CRAC/CLIP data analysis.

## STAR Methods

### fCRAC procedure

The detailed, stepwise fCRAC protocol and associated information is provided in the **Supplemental File**.

### Experimental model and study participant details

#### HeLa cell culture

HeLa cell lines (female) were obtained from DSMZ (Cat#ACC57) and maintained between passages 3 and 35. Cells were grown at 37 °C under 5% CO_2_ in RPMI Glutamax medium (Gibco; Cat#61870036) supplemented with 10% FBS, 1 mM pyruvate sodium, 1% MEM non-essential amino acids and 100 µg/mL penicillin/streptomycin. Cultures were routinely screened for Mycoplasma contamination using the MycoAlert Assay (Lonza; Cat#LT07). Cell line identity was not further verified.

#### K562 cell culture

K562 cells (female; ATCC; Cat#CCL-243; cell identify not further verified) were cultured at 37 °C in RPMI 1640 Medium with GlutaMAX further supplemented with 1x final concentration Antibiotic-Antimycotic (Gibco; Cat#15240096) and 10% fetal calf serum (Sigma; Cat#F2442). Cells were grown at 37 °C under 5% CO_2_ to a density of 0.5–1 × 10^6^ cells/mL, then diluted or used for experiments.

#### HEK293 Cell culture

Flp-In T-REx cells (female; Thermo Fisher; Ca#R78007; cell identify not further verifieds) were cultured at 37°C with 5% CO 2 in DMEM (Thermo-Fisher; Cat#0566016) supplemented with 10% tetracycline-tested FBS (Sigma; Cat#F2442) before transfection as described below.

### Method Details

#### Generation of HeLa cells expressing tagged Cyclon

The human CYCLON sequence (NM_024098.4, CCDS7993.1) was synthesized as a G-block (IDT) and cloned by Gibson assembly into the Nhe1- and Pac1-digested lentiviral pSico backbone together with a C-terminal HIS8-Ala4-FLAG tag amplified from the positive control plasmid (pcDNA5-FRT-TO-Nprot-HF_GC3opt, Addgene #157732). All constructs were validated by Sanger sequencing. HeLa (DSMZ) cells were infected with the resulting viral particles and sorted using a 100 µm nozzle based on GFP-actin expression included in the vector (Aria Ilu, BD Biosciences). CYCLON expression and localization were verified by immunofluorescence using primary (Anti-FLAG M2 monoclonal antibody (mouse); Sigma-Aldrich; Cat#F1804) and secondary (Goat anti-Mouse IgG Secondary Antibody AF546; Invitrogen; Cat#A-11030) antibodies.

#### Generation of positive control cells

As a positive control for the fCRAC procedure, HEK-293FT cells expressing HF-tagged N protein from SARS-CoV-2 virus were generated^10^. Cells were grown in 15 cm dishes in antibiotic-free media to 70% of confluence on the day of transfection, then transfected with plasmid encoding HF-tagged N protein (pcDNA5-FRT-TO-Nprot-HF_GC3opt; Addgene; Cat#157732). Transfection mix contained 25 µg of plasmid, 90 µl of Lipofectamine 2000 (Thermo-Fisher; Cat#11668030) and 3 ml of OptiMEM (Gibco; Cat#10149832) following the Lipofectamine protocol. After 4 hours, media was replaced with fresh media containing 1 ug/mL of doxycycline. Cells were cultured for 48 hours before cross-linking, following the fCRAC protocol (see **Supplemental File**).

#### Immunofluorescence

HeLa cells plated on 12 mm coverslips were fixed in 4% paraformaldehyde for 5 minutes, permeabilized with 0.5% (v/v) Triton X-100 in PBS for 5 minutes, then blocked with 10% Bovine Serum Albumin (in 0.1% Tween PBS (10% BSA-PBST) for 1 hour. Cells were incubated with anti-FLAG primary antibody (Sigma; Cat#F1804), 1/100 dilution in 3% BSA-PBST) overnight at 4°C. Then, cells were washed 3 times in 3% BSA-PBST, incubated with Goat anti-Mouse IgG Secondary Antibody AF546 (Invitrogen; Cat#A-11030) for 1 hour at room temperature and washed 3 times in 3% BSA-PBST. Finally, cells were stained with Hoechst (Thermo; Cat# H1399; 1:10000 dilution in PBS) for 3 minutes, and mounted on a coverslip using Mowiol solution (Sigma; Cat# 81381). Images were acquired using the Zeiss AxioImager M2 using the Plan-Apochromat 40x/0.95 lens.

#### Structural modeling

Human nuclear pre-60S ribosomal subunit (State I2; PDB:8FL3^33^) was visualized and modified using ChimeraX (version 1.11). Contacts were identified using an overlap of -0.4.

### Quantification and Statistical Analysis

All quantification and statistical analysis was performed using trxtools, with tests and interpretation as described in the Figure legends. All experiments were performed at least twice, and no statistical methods were used to estimate sample size.

## Key Resources Table

| REAGENT or RESOURCE | SOURCE | IDENTIFIER |
| --- | --- | --- |
| Antibodies |  |  |
| Anti-FLAG M2 monoclonal antibody (mouse) | Sigma-Aldrich | Cat#F1804 |
| Goat anti-Mouse IgG Secondary Antibody, AF546 | Invitrogen | Cat#A-11030 |
| IRDye 680RD Goat Anti-Mouse IgG | Licor | Cat#926-68070 |
| Chemicals, peptides, and recombinant proteins |  |  |
| 3x FLAG peptide | Sigma | Cat#F4799 |
| Experimental models: Cell lines |  |  |
| HeLa cell line | DSMZ | Cat#ACC57 |
| HEK293 Flp-In T-REx cell line | Thermo Fisher | Cat#R78007 |
| K562 cell line | ATCC | Cat#CCL-243 |
| Oligonucleotides |  |  |
| See Table S1 |  |  |
| Recombinant DNA |  |  |
| Human CYCLON sequence G-block | IDT | Sequence: NM_024098.4 |
| SARS-CoV-2 N protein plasmid: pcDNA5-FRT-TO-Nprot-HF_GC3opt | Addgene | Cat#157732 |
| Software and algorithms |  |  |
| Trxtools | GitHub | Repo: github.com/TurowskiLab/trxtools |
| Trxtools Example Analysis | GitHub | Repo: github.com/TurowskiLab/example-trxtools-Cyclon/ |

## Supplemental File

### Supplemental File Contents

1. fCRAC reagent and equipment list
2. fCRAC buffers
3. fCRAC procedure
4. fCRAC oligonucleotides and linkers
5. Trxtools standardised directory structure
6. Trxtools file naming conventions
7. Supplemental references

## Supporting information

Supplemental File (Protocol and trxtools intro)

## Acknowledgements

We thank the Microcell core facility of the Institute for Advanced Biosciences (UGA - Inserm U1209 - CNRS 5309), especially Mylene Pezet and Solenne Dufour, for their assistance with the Aria Ilu (BD, Biosciences) equipment. This facility belongs to the IBISA-ISdV platform, a member of the national infrastructure France-BioImaging supported by the French National Research Agency (ANR-10-INBS-04). TWT was supported by the Polish National Agency for Academic Exchange (PPN/PPO/2020/2/00004/U/00001), TWT and JM were supported by National Science Center (2020/39/D/NZ2/02115 and 2023/51/I/NZ2/02200). JM was supported by EMBO (SEG #10959) and Institute of Biochemistry and Biophysics Intramural grant FBW-SD-03/2024.NR was supported by a Scottish Government Chief Scientist Office/NHS Education for Scotland Clinical Lectureship (PCL/23/07) and an Academy of Medical Sciences’ Starter Grant for Clinical Lecturers award (SGCL033\1114). AGS was supported by an EMBO Scientific Exchange Grant (11196). DT, MM and AH were supported by Wellcome Principal Research Fellowships to DT [109916, 222516].

## Author contributions

AH, TWT, NR, DT: method development

JM, TWT: Code development

NR, AGS, AE, DT, AH, TWT: designed experiments

NR, AGS, MM: performed experiments

JM, TWT: Data analysis

All: Data interpretation

All: drafted/revised report

---

**Supplemental Figure 1:**
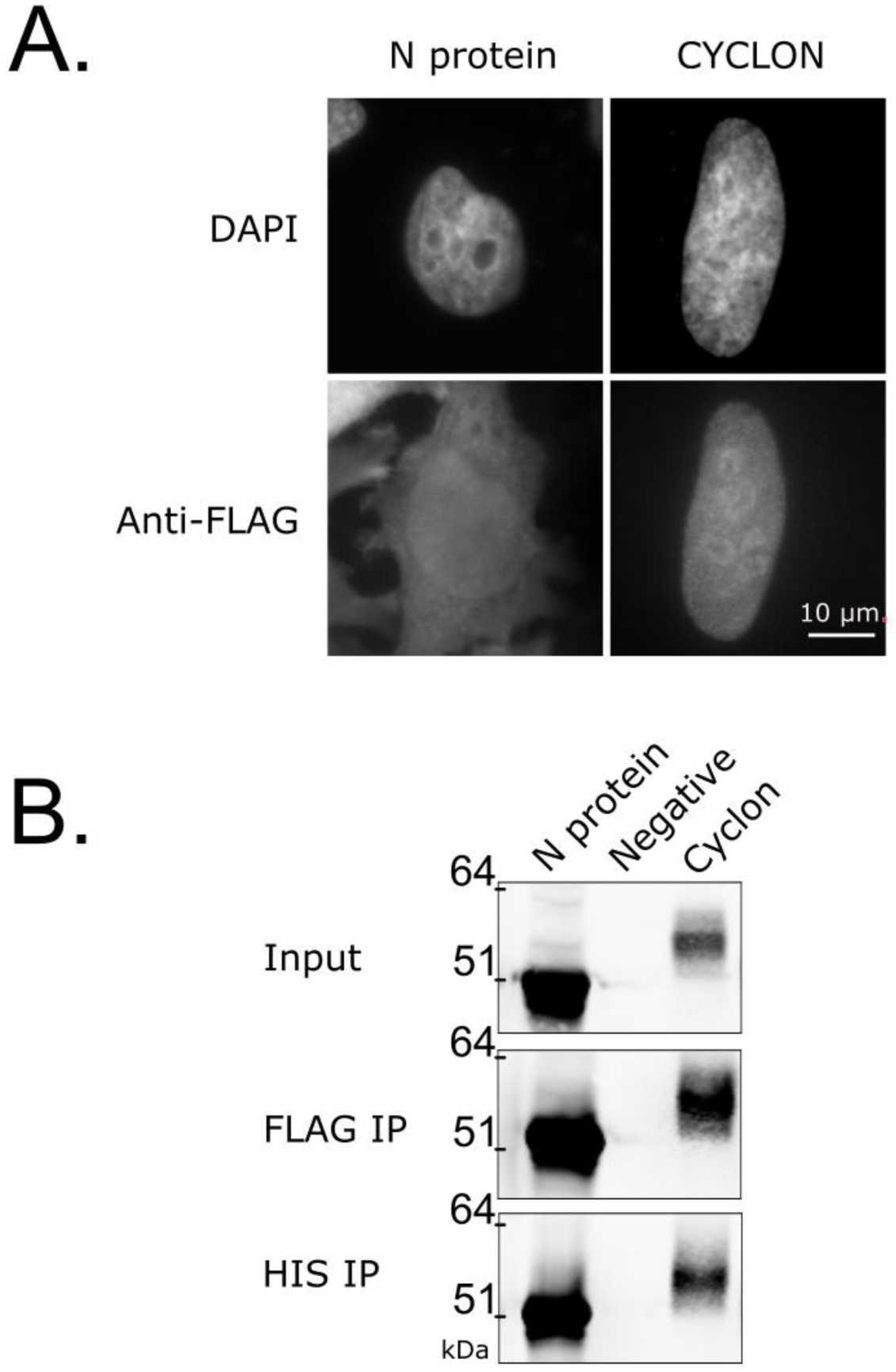
Validation of FLAG-HIS tagged CYCLON protein. **(A)** Immunofluorescence using anti-FLAG antibody and DAPI for FH-tagged CYCLON (CYCLON-HF) protein and the positive control for fCRAC, N protein from SARS-CoV-2. Scale bar indicates 10 µm. **(B)** Immunoblotting for CYCLON protein in total cell lysate (Input) and following sequential anti-FLAG (FLAP IP) and nickel bead (HIS IP) purification.

**Supplemental Figure 2:**
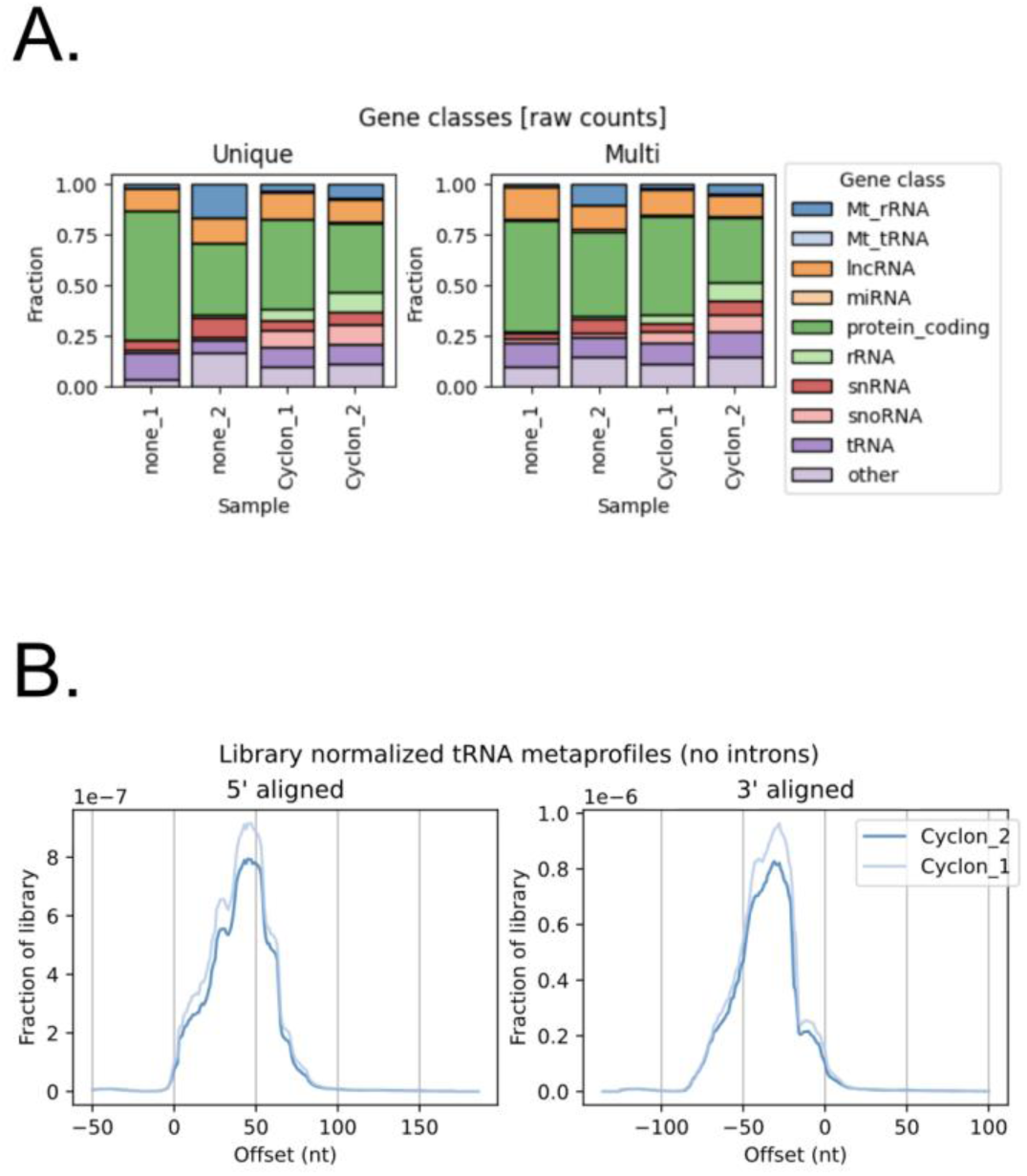
tRNA hits in CYCLON fCRAC. **(A)** Gene classes determined by raw counts (e.g, not TPM-normalized) for CYCLON fCRAC, for both unique and multi-mapping reads. “None” is negative control untagged cell line: in fCRAC, the read distribution of the negative control normally mirrors the real samples, but with much fewer reads, due to cross-contamination during processing. The fraction of tRNA reads in the CYCLON samples was ∼10-12%. **(B)** Metaprofiles for tRNA reads from two replicates of CYCLON fCRAC, aligned to either the 5’ or 3’ end of the gene, demonstrating that a fraction of the hits are from pre-tRNA with unprocessed 5’ or 3’ leaders.

