## Supplemental File (Protocol and trxtools intro) for "Isotope-Free Mapping of protein:RNA Interactions Using fCRAC and trxtools"

Robertson *et al,* 2026

**Supplemental File**

**Contents:**

1. fCRAC reagent and equipment list
2. fCRAC buffers
3. fCRAC procedure
4. fCRAC oligonucleotides and linkers *(including Supplemental Table 1)*
5. Trxtools standardized directory structure *(including Supplemental Table 2)*
6. Trxtools file naming conventions
7. Supplemental references

### 1. fCRAC reagent and equipment list

**Oligonucleotides and linkers:** see *Section 4: fCRAC oligonucleotides and linkers*

**Reagents to prepare 3' linker:**
5′ DNA Adenylation Kit (NEB; cat. E2610L)
IRdye-800CW-DBCO (LiCor; cat. 929-50000)
QIAquick Nucleotide Removal Kit (Qiagen; cat. 28304)

**Plasmid**
Positive control for fCRAC: vector for expression of SARS-CoV-2 N protein.
pcDNA5-FRT-TO-Nprot-HF_GC3opt (Addgene; plasmid #157732[1])

**Buffers components:**
3M Sodium Acetate Solution, pH 5.2
1 M Tris-HCl pH 7.5
1 M Bis-Tris pH 6.5
0.5 M HEPES/KOH pH 7.6
3 M NaCl
5 M LiCl
1 M $MgCl_2$
1 M Imidazole
0.5 M EDTA pH 8
10% IGEPAL CA-630 (Sigma-Aldrich, cat. I8896-50ML)
10% Triton X-100
10% Sodium deoxycholate
10% Sodium dodecyl sulfate
Guanidine hydrochloride (powder)

Urea (powder)
100% ethanol
70% ethanol
80% acetone
Phenol, pH 8
Chloroform
isoamyl alcohol
2-Mercaptoethanol

**Consumables:**
1.5 ml microcentrifuge tubes
5 ml microcentrifuge tubes
5 ml syringes
0.22 µM filter unit for syringe filtration (Millipore; cat. SLMP025SS)
SigmaPrep spin columns (Sigma; cat. SC1000)

**Reagents for protein purification:**
1 mm Zirconia lysis beads (Thistle Scientific, cat. BSP-11079110zx)
EDTA-free protease inhibitor cocktail tablets (Roche cOmplete, cat. 11873580001)
3x FLAG peptide (Sigma; cat. F4799)
Nickel beads: Ni-NTA Agarose (Qiagen; cat. 30210)
anti-FLAG M2 magnetic beads (Milipore, cat. M8823)

**Enzymes and related consumables:**
Lipofectamine 2000 (Invitrogen 11668019)
10 mM ATP (Invitrogen; cat. AM8110G)
GlycoBlue co-precipitant (Invitrogen; cat. AM9516)
RNace-IT Ribonuclease Cocktail (10 U/µL; Agilent; cat. 400720)
RNasin Ribonuclease Inhibitor (Promega; cat. N2511)
T4 Polynucleotide Kinase (NEB; cat. M0201L)
T4 RNA ligase 1 (NEB; cat. M0204L)
Truncated T4 RNA ligase II K227Q (NEB; cat. M0373L)
Proteinase K 20 mg/ml solution (Invitrogen; cat. 25530049)
Superscript IV reverse transcriptase (Invitrogen; cat. 18090010)
Exonuclease I (Exo1; NEB; cat. M0293S)
Phusion High-Fidelity DNA Polymerase (2 U/µl; Thermo Scientific; cat. F530S)
dNTP mixture 2.5 mM of each (e.g. Invitrogen; cat. R72501)

**PAGE reagents:**
4x NuPage LDS sample buffer (Invitrogen; cat. NP0007)
NuPage 4-12% polyacrylamide gel (10 well, 1.5 mm; Invitrogen; cat. NP0321)
20x NuPAGE MOPS SDS Running Buffer (Invitrogen; cat. NP0001)
20x NuPAGE transfer buffer (Invitrogen; cat. NP0006)

Amersham Protran Nitrocellulose Membrane (Cytiva, cat. 15209814)
Primary antibody for detecting Flag-tagged proteins: anti-M2-FLAG antibody (Sigma-Aldrich; cat. F1804)
Secondary antibody: IRDye 680RD Goat Anti-Mouse IgG (Licor, 926-68070)

**Library purification consumables:**
50 bp DNA ladder (Neb; cat. N3236S)
Metaphor agarose (Lonza; cat. 50180)
Silica-column based PCR product purification kit (for example, DNA Clean & Concentrator-5; Zymo Research; cat. D4013)
Silica-column based agarose gel purification kit (for example, Zymoclean Gel DNA Recovery Kit; Zymo Research; cat. D4007)

**Equipment:**
Amersham Life Science Ultraviolet Crosslinker, or equivalent device capable of UVC cross-linking
Cell scrapers (Thermo Scientific, cat. 179707PK)
Magnetic rack for 1.5 ml microcentrifuge tubes
Rotating wheel for 1.5 ml and 5 ml microcentrifuge tubes
Licor Odyssey M Model 3350, or other device capable of reading fluorescence on nitrocellulose membrane and giving 1:1 scale printout as cutting guide

#### 2. fCRAC buffers

**RNace-IT cocktail:** dilute 1:10 to 1 U/µl and store aliquots at -80 degrees

**Buffer LB:** 50 mM Tris-HCL pH 7.5, 0.1 M NaCl, 5 mM $MgCl_2$, 1% IGEPAL CA-630, 0.5% sodium deoxycholate, 0.1% sodium dodecyl sulfate

**Buffer C:** 50 mM Tris-HCL pH 7.5, 50 mM NaCl, 0.1% IGEPAL CA-630

**Buffer FA2:** 50 mM HEPES/KOH pH 7.6, 500 mM NaCl, 1 mM EDTA, 1% Triton X-100, 0.1% sodium deoxycholate

**Buffer FA3:** 10 mM Tris-HCL pH 7.5, 250 mM LiCl, 1 mM EDTA, 0.5% IGEPAL CA-630, 0.5% sodium deoxycholate

**Buffer WB1:** 50 mM Tris-HCL pH 7.5, 500 mM NaCl, 0.1% IGEPAL CA-630, 10 mM imidazole, 6 M guanidine hydrochloride

**Buffer WB2:** 50 mM Tris-HCL pH 7.5, 500 mM NaCl, 0.1% Triton X-100

**Buffer WB3:** 50 mM Tris-HCL pH 7.5, 500 mM NaCl, 0.1% Triton X-100, 8 M urea

**10x reaction buffer R1:** 700 mM Tris-HCL pH 7.5, 100 mM $MgCl_2$

**10x reaction buffer R2:** 700 mM Bis-Tris pH 6.5, 100 mM $MgCl_2$ (aliquots can be frozen at -20 degrees)

**Buffer BE (elution buffer): Buffer WB1** with an additional 300 mM Imidazole. To make, combine 3.5 ml of **Buffer WB1** with 1.5 ml of **1 M imidazole.**

**Buffer PK (proteinase K): Buffer C** with additional 5 mM EDTA and 1% SDS. To make, combine 9 ml of **buffer C** with 0.1 ml of 0.5 M EDTA and 1 ml of 10% SDS.

**Buffer PCI:** phenol-chloroform-isoamyl alcohol mixture in ratio 25:24:1 (phenol pH 8)

#### 3. fCRAC procedure

**Cell growth and cross-linking (adherent cells)**

1. Grow cells to approximately 70% of confluency (Note: ribosome synthesis machinery may begin to shut down at higher confluency)
2. Aspirate media and wash cells with warm PBS, being sure to aspirate all to leave dry layer of cells
3. Place plate containing cells in container of ice (on the ice bed). Take lid off plate.
4. Irradiate cells with UVC (254 nm) to a total dose of 400 mJ/cm2
5. Add 5 ml of cold PBS to the dish and detach the cells by using a cell scraper
6. Transfer cell suspension liquid to a 15 ml tube and centrifuge (4°C, 350g, 5 min)
7. Aspirate PBS and store cell pellet at -80°C until ready for use

*Cells grown in suspension, such as K562, can be cross-linked in their full media, using a Vari-X-Linker crosslinking device*[2] *and quartz dish. We have previously published a protocol describing this approach*[3].

**Cell Lysis**

1. For each experimental sample, resuspend pellets containing up to 50 million cells in 3 ml of ice-cold **buffer LB**, supplemented with 1 protease-inhibitor cocktail tablet per 50 ml of buffer
2. Incubate on ice for 10 min

3. Add 0.5 ml of Zirconia lysis beads and vortex tube on a table-top vortex on the maximum setting for 30 sec
4. Remove the insoluble fraction: pre-clear lysate by centrifugation (4°C, 12,000 g, 5 min)
5. Load the supernatant lysate into 5 ml syringe and pass filter through a 0.22 µm PES filter syringe unit into a 5 ml microtube

**Anti-Flag immunoprecipitation**

*We recommend taking quality control (QC) samples to follow the protein through the purification steps. Suggested collection points are indicated in italics throughout the protocol.*
*QC - collect input sample: take 30 µl (1%) of lysate into new tube, and freeze*

1. Wash 100 µl of anti-FLAG M2 magnetic beads per sample: aliquot total amount of beads requirement into 1.5 ml tube and use magnetic rack to wash twice with 1 ml of **buffer LB**, and resuspend back in original volume
2. Add 100 µl of washed beads to each lysate sample
3. Incubate on a rotating wheel for 2 h at 4°C
4. Transfer beads to one 1.5 ml tube per sample - place the 1.5 ml tube in the magnetic rack, transfer 1 ml of lysate and discard lysate once beads settled, and repeat until all beads are transferred

*QC - collect Flag flow-through sample: before discarding first batch of lysate, take 30 µl of the supernatant lysate into a new tube, and freeze.*

5. Wash beads 3 times with 1 ml of **buffer LB**
6. Wash beads 2 times with 1 ml of **buffer C**
7. Resuspend beads in 300 µl of **buffer C**

**RNA footprinting**

1. Prepare RNase digestion RNAce-IT solution: per sample mix 300 µl of **buffer C,** and 0.5 units of RNace-IT

*This is for a "higher digestion". Depending on the extent that the protein protects the cross-linked RNA, it may be necessary to do a lower digestion, in which case use 0.04 units of RNace-IT per sample, again in 300 µl of* ***buffer C***

2. Add 300 µL of the RNAce-IT solution to each lysate the sample and incubate for 10 min in a heat-block set to 23°C, with shaking at 1000 rpm.
3. Transfer tubes to the magnetic rack and remove solution from beads
4. Wash beads 1 times with 1 ml of **buffer LB**
5. Wash beads 3 times with 1 ml of **buffer FA2 -** place tubes in ice bucket and agitate gently for 3 min before placing back on magnetic rack
6. Wash beads 3 times with 1 ml of **buffer FA3 -** agitate on ice as before
7. Wash beads 2 times with 1 ml of **buffer C**

**Elution from anti-Flag beads**

1. Remove tubes from the magnetic rack
2. Resuspend beads in 200 µl of **buffer LB,** then add 6 µl of 3xFLAG peptide (5 mg/ml, 150 µg/mL final concentration).
3. Incubate tubes the samples for 5 min in a heat block set 37°C, with shaking at 1200 rpm
4. Place the samples tubes on the magnetic rack and collect eluate into the new 1.5 ml tube
5. Repeat elution in further 200 µl of **buffer LB** supplemented with 6 µl of 3xFLAG peptide
6. Combine the eluates into one 1.5 ml tube

*QC - collect Flag-elution sample: take 16 µl of eluate into new tube, and freeze*

**His-tag purification**

1. Add 800 µl of **buffer WB1** to 400 µl the eluate to denature the proteins
2. Prepare Ni-NTA beads: per sample, wash 75 µl of nickel beads twice with **buffer WB1.** To do this, add 1 ml of **buffer WB1** to the bead volume**,** centrifuge at 1000 g for 30 sec and discard supernatant. After last wash, resuspend beads back in original volume
3. Add nickel beads to the eluates
4. Incubate on a rotating wheel for 2 h at room temperature
5. Wash beads 3 times with 1 ml of **buffer WB1,** following procedure in step 2
6. Wash beads 2 times with 1 ml of **buffer C**
7. Resuspend beads in with 600 µl of **buffer C**

*QC - collect nickel bead sample: take 12 µl of bead suspension, and freeze*

**Dephosphorylation**

1. Load remaining bead solution onto SigmaPrep spin column, with column placed in a collection tube. *From now on all the enzymatic and washing steps will be performed on the column*
2. Wash the beads 2 times with 0.75 ml of 50 mM Bis-Tris, pH 6.5: For washes on column, add the buffer directly on the beads, incubate at room temperature for 30 sec, then remove all the spin buffer into collecting tube by centrifugation (10 sec, room temperature, 300 g). Close the bottom of the tube with the cap to keep the mixture in the column during incubations
3. Prepare reaction mix: per sample, mix 8 µl of **10x buffer R2,** 4 µl of T4 Polynucleotide Kinase, 2 µl of RNasin ribonuclease inhibitor, and 64 µl of distilled water (MQ). Ensure beads on columns are dry (e.g. previous wash buffer spun out) then place cap on bottom of column
4. Add 80 µl of reaction mixture onto columns, and incubate in heat block for 30 min at 37°C

5. Move the columns to room temperature, open the lids of the columns, then remove the caps and place the columns back into the collection tubes
6. Wash beads 1 x with 0.5 ml of **buffer WB1**
7. Wash columns 3 x with 0.75 ml of **buffer C.** Make sure to remove any remaining wash buffer
8. Close the bottom of the column with the cap.

**3’ Linker ligation**

1. To each sample add 80 µl of prepared reaction mix: per sample, mix 8 µl of **10x buffer R1,** 4 µl of 20 µM fluorescent 3’ linker, 4 µl of truncated T4 RNA ligase II K227Q, 2 µl of RNasin ribonuclease inhibitor, 42 µl of MQ, and 20 µl of PEG8000 (provided with the ligase). Ensure beads on columns are dry (e.g. previous wash buffer spun out) then place cap on bottom of column
2. Add 80 µl of reaction mix to columns and incubate the samples in a heat block for overnight at 16°C, with mixing (e.g. 500 rpm).
3. Move the columns to room temperature, open the lids of the columns, remove the caps and place the columns back into the collection tubes
4. Wash the columns:
    a. 1 x 0.5 ml of **buffer WB1**
    b. 1 x 0.75 ml of **buffer WB2**
    c. 1 x 0.5 ml of **buffer WB3**
    d. 1 x 0.75 ml of **buffer WB2**
    e. 4 x 0.75 ml of **buffer C**

**Buffer exchange washes**

A buffer exchange wash is used to remove unligated linker. After removing the cap, wash columns:

1. 1x with 0.5 ml of **buffer WB1**
2. 1x with 0.75 ml of **buffer WB2**
3. 1x with 0.5 ml of **buffer WB3**
4. 1x with  0.75 ml of **buffer WB2**
5. 4x with 0.75 ml of **buffer C**

**PNK treatment**

1. Prepare reaction mix: per sample, mix 8 µl of **10x buffer R1,** 8 µl of 10 mM ATP, 2 µl of T4 Polynucleotide Kinase, 2 µl of RNasin ribonuclease inhibitor, 40 µl of MQ, and 20 µl of PEG8000 (provided with ligase enzyme)
2. Ensure beads on columns are dry (e.g. previous wash buffer spun out) then place cap on bottom of column
3. Incubate in a heat block set to 37°C for 30 min

**5’ linker ligation**

1. To the reaction mixture already on each column, directly add 4 μl of T4 RNAase ligase 1 and 2 μl of 5’ barcoded linker (100 μM). Use a different barcoded linker for each sample.
2. Incubate on a heatblock set to 16°C for 3 h, then for a further 2 h at 25°C, with shaking (500 rpm)
3. Remove cap, and wash columns 3 times with **buffer WB1**

**Elution from nickel beads and precipitation**

1. Ensure beads on columns are dry (e.g. previous wash buffer spun out) then place cap on bottom of column
2. Resuspend beads in 50 μl of **buffer BE** and incubate on a heatblock set at 23°C with mixing (800 rpm) for 5 min
3. Remove caps and centrifuge eluate into 1.5 ml microcentrifuge tube (30 sec, room temperature, 300 g)
4. Repeat steps 1-3 above for a second elution, collecting eluate into the same tube
5. To precipitate RNA:protein complexes, add 3 μl of GlycoBlue and 900 μl of 100% ethanol, mix well, and precipitate overnight at -20°C

**PAGE gel and transfer**

1. Centrifuge to collect precipitated samples (20 min, 4°C, 12,000 g)
2. Carefully remove supernatant from pellet, and add 1 ml of cold 80% acetone
3. Vortex for 15 sec, then centrifuge (5 min, 4°C, 12,000 g)
4. Repeat steps 2-3 for a second wash
5. Remove all acetone and let pellet air dry (normally 2-3 min - avoid overdying as this will make the pellet harder to resuspend)
6. Add 12 μl of MQ to the pellet and allow to rehydrate at room temperature for 10 min
7. Pipette sample up and down several times to resuspend, then add 4 μl of 4x NuPage LDS sample buffer, together with 2-Mercaptoethanol to a final concentration of 2%. Mix well by pipetting
8. Heat denature samples by placing on a heat-block set to 65°C for 10 min
9. Load samples onto a NuPage 4-12% gel. Load a protein ladder in the first well, and leave 1 empty well between each sample
10. Run gels at 130 V in 1x MOPS buffer, until the ladder is fully extended. Keep gel tank cool during run by placing in ice-filled box
11. Transfer gel onto a nitrocellulose membrane in 1x transfer buffer supplemented with 10% methanol, running at 100 V for 2 h. Keep transfer tank cool during run
12. Image the membrane, and print out 1:1. The labelled, cross-linked RNA should be visible as a smear above the expected size of the protein
13. Annotate the smear of RNA:protein complexes using a marker pen on the printout

14. Place this cutting guide under a flat piece of glass, and place the membrane on the glass with a piece of clingfilm beneath. Using a clean, sharp disposable scalpel, cut out the region of interest
15. Cut the excised band into small squares (about 2 mm diameter) and place in a 1.5 ml microcentrifuge tube. Be sure not to transfer any clingfilm remnants.

**Proteinase K treatment**

4. To excised bands, add 500 µl **Buffer PK** and 5 µl of proteinase K (20 mg/ml; 100 µg total)
5. Incubate in a heatblock set to 55°C for 1 h, with shaking at 1400 rpm
6. Add 50 µl of 3M sodium acetate pH 5.2
7. In a fume hood, carefully add 500 µl of **buffer PCI** and vortex tubes for 30 sec at room temperature

***Warning:*** *phenol-containing solutions are hazardous. Handle in a fume hood and comply with locally-agreed safety procedures*

5. Centrifuge at 16,000 g for 5 min at 4°C
6. Transfer 350 µl of upper phase into new 1.5 ml microcentrifuge tube and add 2 µl of Glycoblue coprecipitant and 1 ml of 100% ethanol
7. Precipitate overnight at -20 degrees

**Reverse transcription and primer removal**

1. Centrifuge tubes at 16,000 g for 20 min at 4 degrees C.
2. Carefully remove the supernatant and discard
3. Wash the pellet by adding 1 ml of 70% ethanol and vortexing
4. Centrifuge tube at 16,000 g for 5 min at 4°C
5. Carefully remove the supernatant and discard, and allow pellet to air dry
6. Resuspend the pellet by pipetting up and down in a 13 µl of the following mixture: 1 µl of fCRAC RT oligo (10 µM), 4 µl of 2.5 mM dNTPs, and 8 µl ul of MQ
7. Incubate for 3 min in a heatblock set to 80°C
8. Place tube on ice for 5 min
9. Briefly centrifuge in a table-top centrifuge to collect condensate
10. Add 10 µl of the following mixture: 4 µl of 5x RT buffer, 1 µl of 100 mM DTT (both provided with Superscript IV), 1 µl of RNasin RNAse inhibitor, and 1 µl of Superscript IV enzyme
11. Incubate in a heatblock set to 50°C for 15 min
12. Incubate at room temperature for 5 min
13. Incubate on ice for 3 min
14. A 2 µl of Exo1 enzyme and incubate for 30 min at 37°C
15. Incubate at 80°C for 20 min to inactivate Exo1

**Library amplification**

1. For each sample, prepare a 50 µl PCR reaction comprising 10 µl of 5X Phusion HF buffer (provided with polymerase), 5 µl of 2.5 mM dNTP, 1 µl of **primer P5** (at 10 µM), 1 µl of **primer BC01** (at 10 µM), 0.3 µl of Phusion HF polymerase, and 30.7 µl of MQ. Add 2 µl of cDNA
2. Run PCR program: 98°C for 30 sec, followed by 21 cycles of 98°C for 10 s, 65°C for 30 sec and 72°C for 30 sec, followed by a final extension at 72°C for 5 min
3. Clean up PCR using silica-column based PCR kit, and elute in 10 µl of MQ.
4. Mix PCR eluate with loading dye, and run on 3% Metaphor gel including a DNA stain (e.g. SYBR safe). Run a 50 bp ladder for scale
5. Image the gel and print out the image at 1:1 scale
6. Using the print out as a cutting guide, excise agarose containing fCRAC library from gel using a clean, sharp scalpel
7. Extract library from excised gel bands using silica-column based extraction kit, and elute in 20 µl of MQ
8. Quantify eluted library

*For sequencing, about 5-50 ng of purified DNA is required. To generate the final library for sequencing, adjust the number of PCR reactions per sample and the number of PCR cycles, aiming to achieve total product in this range with minimum number of PCR amplification steps. For example, for each fCRAC sample, perform 3x 50 µl reactions at 18 PCR cycles, and pool during column purification.*

**QC: Western blots to follow protein purification**

We recommend running the four quality control samples for the protein purification on a Western Blot (*Input, Flag flow-through, Flag eluate and Nickel beads*):

1. For the Input, Flag flow-through and Flag eluate samples: add 4x loading buffer to 1x final concentration, boil for 5 min, and load and run on gel as in *PAGE gel and transfer* section of main protocol

*The nickel bead sample is in denaturing buffer and needs to be precipitated before further processing:*

1. Add 3 µl of Glycoblue and 9 volumes (108 µl) of ice-cold 100% ethanol to the 12 µl sample
2. Allow to precipitate in -80°C freezer for 1 hour (alternatively, overnight at -20°C)
3. Centrifuge at 16,000 g for 20 min at 4°C
4. Carefully remove supernatant
5. Wash pellet with 80% cold acetone and then allow to air dry
6. Add 12 µl of water and allow pellet to rehydrate. Resuspend by pipetting, then add 4 µl of 4X loading buffer
7. Boil and load as for other samples

Probe membranes with anti-flag antibody (Agilent 200474) and appropriate secondary.

### 4. fCRAC oligonucleotides and linkers

**Ordering and processing of fluorescent 3' linker:**

The fluorescent 3' linker was developed from that used in iCLIP[4]. The following oligo was ordered from IDT:

5'-NGCTtcNNNNNNAGATCGGAAGAGCACACGTCTGAAAAAAAAAAAAA/iAzideN/AAAAAAAAAAAAA/3Bio/-3'

The 5' element contains random nucleotides, followed by a region (underlined) complementary to the PCR primers. Downstream there is a polyA tract with an internal azide moiety (iAzide) to facilitate fluorescent coupling. The linker also includes a 3' biotin modification, although this is not utilized in the version of the protocol described here.

Before use, the adaptor was phosphorylated, pre-adenylated and conjugated to the IRDye (IRdye-800CW-DBCO), following the protocol published by Zarnegar *et al.*[4]

***Supplemental Table 1 – fCRAC oligonucleotides and linkers***

| Item | Description | Sequence | |
|---|---|---|---|
| 3' linker | Fluorescent 3' linker | *See note above for sequence and processing information* | |
| 5' linkers | Illumina barcoded 5' linkers | L5Aa | 5′ - invddT-ACACrGrArCrGrCrUrCrUrUrCrCrGrArUrCrUrNrNrNrUrArArGrC-OH - 3′ |
| | | L5Ab | 5′ - invddT-ACACrGrArCrGrCrUrCrUrUrCrCrGrArUrCrUrNrNrNrArUrUrArGrC-OH - 3′ |
| | | L5Ac | 5′ - invddT-ACACrGrArCrGrCrUrCrUrUrCrCrGrArUrCrUrNrNrNrGrCrGrCrArGrC-OH - 3′ |
| | | L5Ad | 5′ - invddT-ACACrGrArCrGrCrUrCrUrUrCrCrGrArUrCrUrNrNrNrCrGrCrUrUrArGrC-OH - 3′ |
| | | L5Ba | 5′ - invddT-ACACrGrArCrGrCrUrCrUrUrCrCrGrArUrCrUrNrNrNrArGrArGrC-OH - 3′ |
| | | L5Bb | 5′ - invddT-ACACrGrArCrGrCrUrCrUrUrCrCrGrArUrCrUrNrNrNrGrUrGrArGrC-OH - 3′ |
| | | L5Bc | 5′ - invddT-ACACrGrArCrGrCrUrCrUrUrCrCrGrArUrCrUrNrNrNrCrArCrUrArGrC-OH - 3′ |
| | | L5Bd | 5′ - invddT-ACACrGrArCrGrCrUrCrUrUrCrCrGrArUrCrUrNrNrNrUrCrUrCrUrArGrC-OH - 3′ |
| | | L5Ca | 5′ - invddT-ACACrGrArCrGrCrUrCrUrUrCrCrGrArUrCrUrNrNrNrCrUrArGrCrN-OH - 3′ |

| | | | |
|---|---|---|---|
| | | L5Cb | 5′ - invddT-ACACrGrArCrGrCrUrCrUrUrCrCrGrArUrCrUrNrNrNrUrGrGrArGrCrN-OH - 3′ |
| | | L5Cc | 5′ - invddT-ACACrGrArCrGrCrUrCrUrUrCrCrGrArUrCrUrNrNrNrArCrUrCrArGrCrN-OH - 3′ |
| | | L5Cd | 5′ - invddT-ACACrGrArCrGrCrUrCrUrUrCrCrGrArUrCrUrNrNrNrGrArCrUrUrArGrCrN-OH -3’ |
| **fCRAC RT primer** | Primer for reverse transcription | 5’-CAGACGTGTGC-3’ | |
| **P5_forward** | Forward PCR primer for library amplification | 5′ - AATGATACGGCGACCACCGAGATCTACACTCTTTCCCTACACGACGCTCTTCCGATCT - 3′ | |
| **BC01 (P7)** | Reverse PCR primer for library amplification | 5’-CAAGCAGAAGACGGCATACGAGATCGTGATGTGACTGGAGTTCAGACGTGTGCTCTTCCGATCT-3’ | |

#### 5. Trxtools standardised directory structure

The accompanying example pipeline are distributed as a ready-to-use package via GitHub. This includes a standardized bioinformatics directory structure and file-naming convention, removing the need for metafiles.

The general concept of a standardized directory is rooted in Cookiecutter Data Science[5], which provides a data science project template. The general structure was adapted to the needs of high-throughput analysis and can be utilized for bioinformatic analysis of high-throughput sequencing projects.

*Supplemental Table 2: Folder structure for standardized bioinformatics analysis*

| Folder | Origin | Comment |
|---|---|---|
| 00_raw | Repository | Action required: place raw FASTQ files here |
| 01_preprocessing | Created by pipeline | Contains preprocessed data |
| 02_alignment | Created by pipeline | Stores alignment outputs |
| 03_FeatureCounts | Created by pipeline | Contains feature count tables. Additional folders contain variants with UMItools deduplication and multi-overlapping read counting enabled. |
| 04_BigWig | Created by pipeline | Holds BigWig files for visualization. Additional folder contains variants with UMItools deduplication. |
| 05_notebooks | Repository | Includes Jupyter notebooks for analysis |
| seq_references | Repository | Action required: reference sequences |
| envs | Repository | Environment files (do not modify) |
| scripts | Repository | Pipeline scripts (do not modify) |
| README.md | Repository | Documentation (do not modify) |

The pipeline generates BigWig files, which can be directly visualized using any genome browser e.g. Integrated Genome Viewer (IGV). Additionally, the template repository provides tools for quality control and data assessment, ensuring robust evaluation of sequencing results. All commands to recreate the example data presented hereare available from the associated repository (*https://*github.com/TurowskiLab/example-trxtools-Cyclon*)*.

#### 6. Trxtools file naming conventions

Input files must follow the name structure:

- adaptor3end_adaptor5end_name.fastq.gz

For clarity in downstream analyses, we recommend that the `name` component is structured as follows:

- AuthorDate_bait_background_condition_replicateNo

Although optional, this convention greatly facilitates data organization and interpretation. The *"AuthorDate"* component indicates the date of the wet-lab experiment. Because the multistep library preparation protocol is often a major source of variability and samples are frequently pooled for sequencing, this identifier provides a unique reference to the library preparation process and helps detect potential experimental anomalies.

The "*bait_background_condition_replicateNo*" component enables flexible filtering and sorting of input data across multiple experiments, including those involving wild-type or mutant baits, multiple bait types, diverse genetic backgrounds, and various treatment conditions. Using underscores to separate fields and maintaining a consistent number of components allows names to be easily parsed into tables with a single command. To ensure this functionality, mutants, complex backgrounds, or treatment options should be separated using a hyphen (e.g., "-").

For example, the first replicate of wildtype Cyclon, the input file was named:

- AG241010_Cyclon_none_120mJ_1

Finally full name of raw input file includes:

- NAGTGGTCNNNNNNAGATCGGAAGAGCACACGTCTG_NNNGTGAGC_AG241010_Cyclon_none_120mJ_1.fastq.gz

This type of naming includes all necessary information, and allows for downstream analysis without other metafiles defining an input. An example of naming for more complex experimental setups is provided in Supplementary Table SX.
